# A biophysical model of viral escape from polyclonal antibodies

**DOI:** 10.1101/2022.09.17.508366

**Authors:** Timothy C. Yu, Zorian T. Thornton, William W. Hannon, William S. DeWitt, Caelan E. Radford, Frederick A. Matsen, Jesse D. Bloom

## Abstract

A challenge in studying viral immune escape is determining how mutations combine to escape polyclonal antibodies, which can potentially target multiple distinct viral epitopes. Here we introduce a biophysical model of this process that partitions the total polyclonal antibody activity by epitope, and then quantifies how each viral mutation affects the antibody activity against each epitope. We develop software that can use deep mutational scanning data to infer these properties for polyclonal antibody mixtures. We validate this software using a computationally simulated deep mutational scanning experiment, and demonstrate that it enables the prediction of escape by arbitrary combinations of mutations. The software described in this paper is available at https://jbloomlab.github.io/polyclonal.

## Introduction

Many viruses evolve antigenically to escape polyclonal antibodies elicited by vaccination or prior infection (Smith *et al*. 2004; Hensley *et al*. 2009; Bedford *et al*. 2014; Eguia *et al*. 2021). Neutralization assays are the gold standard for experimentally assessing if a new viral variant has mutations that erode antibody immunity. However, neutralization assays require generating the actual viral variant(s) for individual testing, and can therefore only be effectively applied retrospectively to a modest number of viral variants of interest (DeGrace *et al*. 2022). For this reason, neutralization assays have difficulty keeping pace with identification of vast numbers of emerging viral variants by genomic epidemiology (Elbe and Buckland-Merrett 2017; Rambaut *et al*. 2020; Viana *et al*. 2022), and so it would be useful to have a method for accurately predicting the antigenic phenotype of viral variants with arbitrary combinations of mutations.

Unfortunately, predicting how a polyclonal serum will neutralize a new viral variant remains a challenge. Deep mutational scanning can systematically measure how large libraries of viral protein variants affect antibody neutralization or binding (Dingens *et al*. 2017; Lee *et al*. 2019; Wu *et al*. 2020; Greaney *et al*. 2021a, 2021c). However, although such high-throughput experimental methods can assess the effects of all single mutants to some viral proteins, the number of possible multiply mutated variants far exceeds the limits of these experiments. Therefore, a variety of computational approaches have been developed that attempt to predict escape by new viral variants. These approaches include basic transformations of deep mutational scanning data (Greaney, Starr and Bloom 2022), models that integrate antigenic data with phylogenetic (Neher *et al*. 2016) or sequence data (Sun *et al*. 2013; Harvey *et al*. 2016), and neural networks that can be trained using deep mutational scanning data (Taft *et al*. 2022) or sequence data alone (Hie *et al*. 2021; Thadani *et al*. 2022).

Here we introduce a new model of viral polyclonal antibody escape that has several advantages over existing computational approaches. Our model is interpretable in terms of underlying biophysical parameters, can be directly fit to experimental deep mutational scanning data, and can predict how new viral variants will be neutralized by the polyclonal sera used to generate the experimental data. We implement our model in a software package and validate it on simulated experimental data. Finally, we demonstrate how the parameters of the model provide quantitative intuition for how mutations combine to escape antibodies that target distinct viral epitopes.

## Results

### The concept of antibody epitopes

Our approach is inspired by the idea that viral antigens can be partitioned into distinct epitopes. This idea can be traced back over four decades to classic experiments on influenza, which tested viral mutants against large panels of monoclonal antibodies (Laver *et al*. 1979; Yewdell, Webster and Gerhard 1979; Webster and Laver 1980). Viral escape mutants were first selected using individual antibodies, and then tested against other antibodies in the panel. A pattern that emerged from these experiments was that groups of antibodies were escaped by similar viral mutants (**Fig. 1**). Antibodies that share common escape mutants were inferred to recognize a common epitope on the viral protein. For instance, in **Fig. 1** antibodies 2 to 5 recognize a similar epitope since they are escaped by many of the same viral mutants.

**Figure 1.**
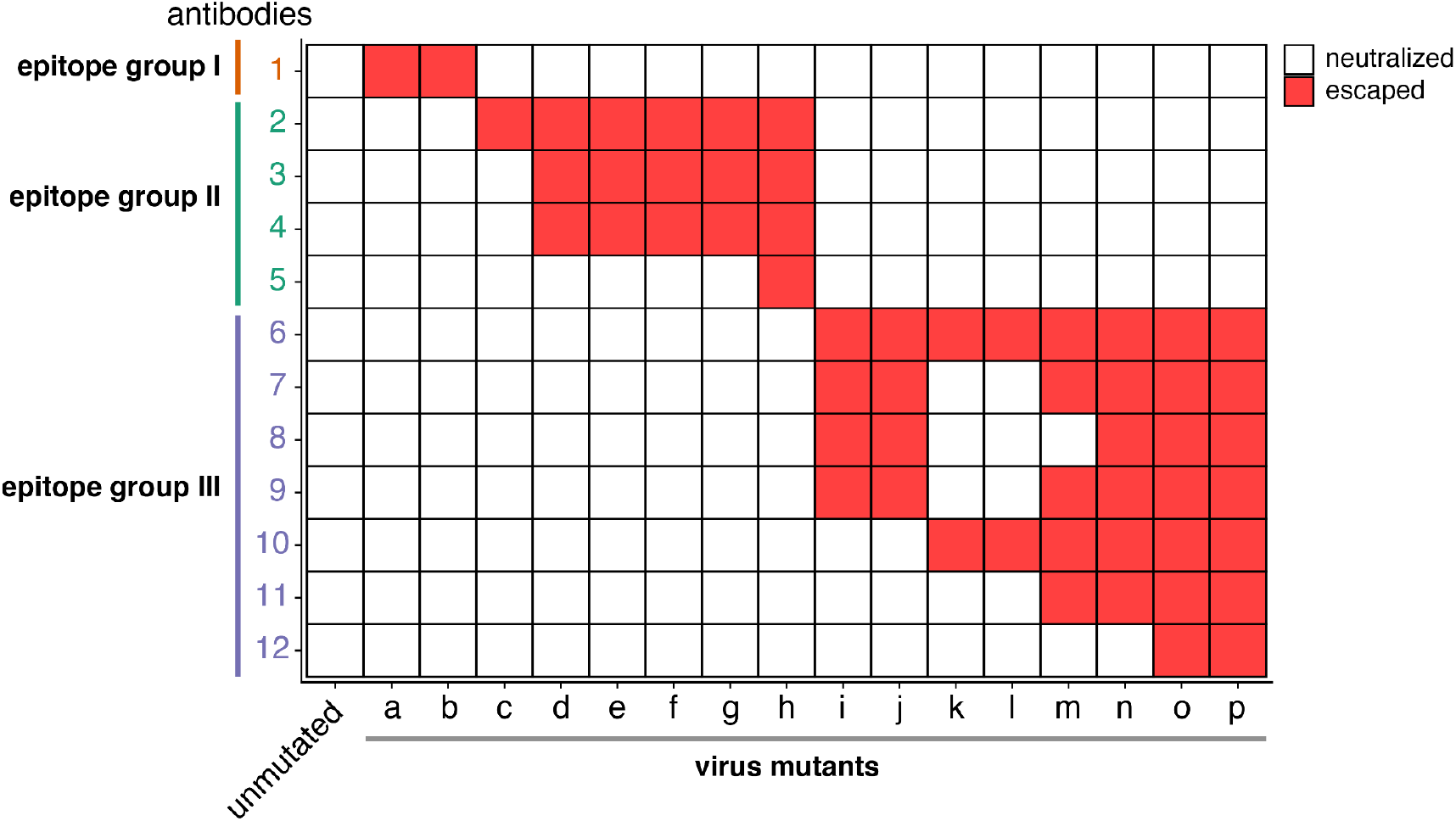
Antibodies can be grouped into epitopes based on whether they share viral escape mutants. Heatmap displaying classic experimental data extracted from Webster and Laver (Webster and Laver 1980). Influenza virus mutants (a-p) were selected using individual antibodies from a large panel of monoclonal antibodies. Each viral mutant was then tested to see if it was neutralized (white) or escaped (red) by the other antibodies in the panel. Antibodies were then grouped into epitopes based on their shared escape mutants.

Based on these studies, the H3 influenza hemagglutinin was divided into five distinct epitopes (Wiley, Wilson and Skehel 1981; Skehel *et al*. 1984). This type of coarse epitope map also provided a straightforward way to conceptualize viral escape from polyclonal serum. Unlike monoclonal antibodies, polyclonal serum can contain multiple antibodies, each recognizing discrete epitopes. As such, any variant that fully escapes polyclonal serum would need to escape antibodies binding at multiple epitopes.

The concept of dividing viral antigens into distinct epitopes has proven to be very useful and continues to be applied to new viruses such as SARS-CoV-2 (Barnes *et al*. 2020; Piccoli *et al*. 2020; Cao *et al*. 2022). However, grouping antibodies by epitopes is a simplifying approximation, as two antibodies targeting a similar epitope may be escaped by slightly different mutations, and some antibodies may bind idiosyncratic regions outside or between major epitopes (Doud, Hensley and Bloom 2017; Piccoli *et al*. 2020; Greaney *et al*. 2021b, 2021c; Starr *et al*. 2021a). For instance, in **Fig. 1** antibodies 6 and 12 are grouped into the same epitope even though some viral mutants only escape one of these two antibodies. Nonetheless, the concept of epitopes provides a valuable way to interpret polyclonal antibody escape and forms the basis of the quantitative approach we take here.

### An epitope-based model of viral escape from multiple antibodies

Given the concept of epitopes described above, consider viral escape from multiple antibodies. For simplicity, consider a hypothetical polyclonal mixture of just two antibodies (1 and 2) that bind distinct epitopes on a viral antigen. The functional activities of these two antibodies in the polyclonal mixture can differ due to antibody-intrinsic factors (e.g., binding affinity and neutralization potency) and extrinsic factors (e.g., antibody concentration). In our hypothetical antibody mixture, we assume antibody 1 has a higher functional activity than antibody 2.

To understand how mutations affect viral escape from this hypothetical two-antibody mixture, consider the viral variants in **Fig. 2**. In the first variant, a single mutation escapes antibody 1, while leaving antibody 2 binding intact. Since antibody 1 has a higher functional activity, losing its contribution causes marked escape from the overall antibody mixture (**Fig. 2A**). However, a viral variant with a single mutation that escapes antibody 2 but leaves antibody 1 binding intact has little escape from the overall antibody mixture, since the more active antibody 1 can still bind (**Fig. 2B**).

**Figure 2.**
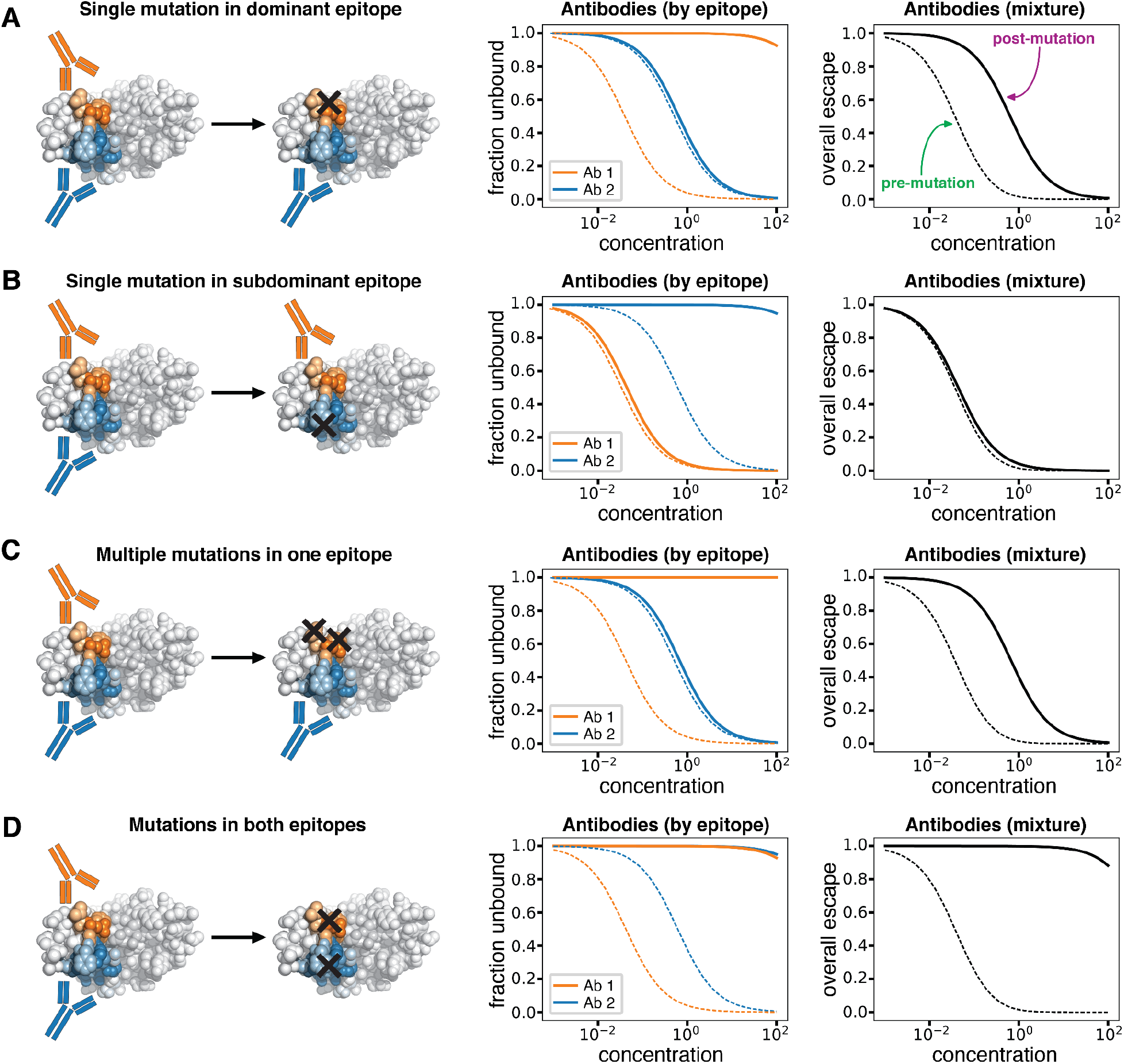
Viral escape from a hypothetical mixture of antibodies targeting two different epitopes. **A)** A single mutation in the dominant (orange) epitope substantially decreases but does not eliminate binding by the antibody mixture. **B)** A single mutation in the subdominant epitope (blue) has minimal effect on binding by the overall antibody mixture since binding of antibodies to the dominant epitope is unaffected. **C)** Multiple mutations in the dominant epitope have an effect that is no greater than a single mutation in this epitope since the remaining binding is due to antibodies targeting the subdominant epitope. **D)** Mutations in both epitopes completely escape binding by the antibody mixture. For each row, the plots show the fraction of antigens unbound by antibody as a function of antibody concentration. The left plot shows the fraction unbound for individual epitopes, and the right shows the overall fraction unbound. The dashed lines indicate binding to the unmutated antigen, while solid lines indicate binding to the mutated antigen.

Importantly, this framework makes distinct predictions about the effects of multiple mutations depending on whether they are in the same or different epitopes. Imagine if the viral variant has two mutations in antibody 1’s epitope. If one of the mutations is already sufficient to mostly escape binding by antibody 1, then the second mutation will have a largely redundant effect (**Fig. 2C**). It follows that multiple escape mutations in the same epitope will have diminishing returns—even when all antibody 1 molecules are unbound, antibody 2 can still bind. However, the situation is very different if the viral variant gains mutations in the epitopes of both antibody 1 and antibody 2. Since both antibodies are escaped, the polyclonal mixture will have no activity (**Fig. 2D**). Note that the principles described above can be readily extended to mixtures of antibodies that target more than two distinct epitopes.

The simple hypothetical example described above is mirrored in real-world data by Kuzmina et al. (Kuzmina *et al*. 2021) on SARS-CoV-2 escape from neutralization by polyclonal serum pooled from vaccinated individuals (**Fig. 3**). The K417N mutation, which is located in the subdominant class 1 epitope (Greaney *et al*. 2021b) of the SARS-CoV-2 receptor binding domain (RBD) has little effect on polyclonal antibody escape. But E484K, which is located in the immunodominant class 2 epitope (Greaney *et al*. 2021b), causes a substantial drop in polyclonal antibody neutralization. But while K417N has no effect on its own, combining K417N with E484K causes a larger drop in neutralization than E484K alone, presumably due to escape from two distinct groups of antibodies in the sera targeting each epitope (**Fig. 3**).

**Figure 3.**
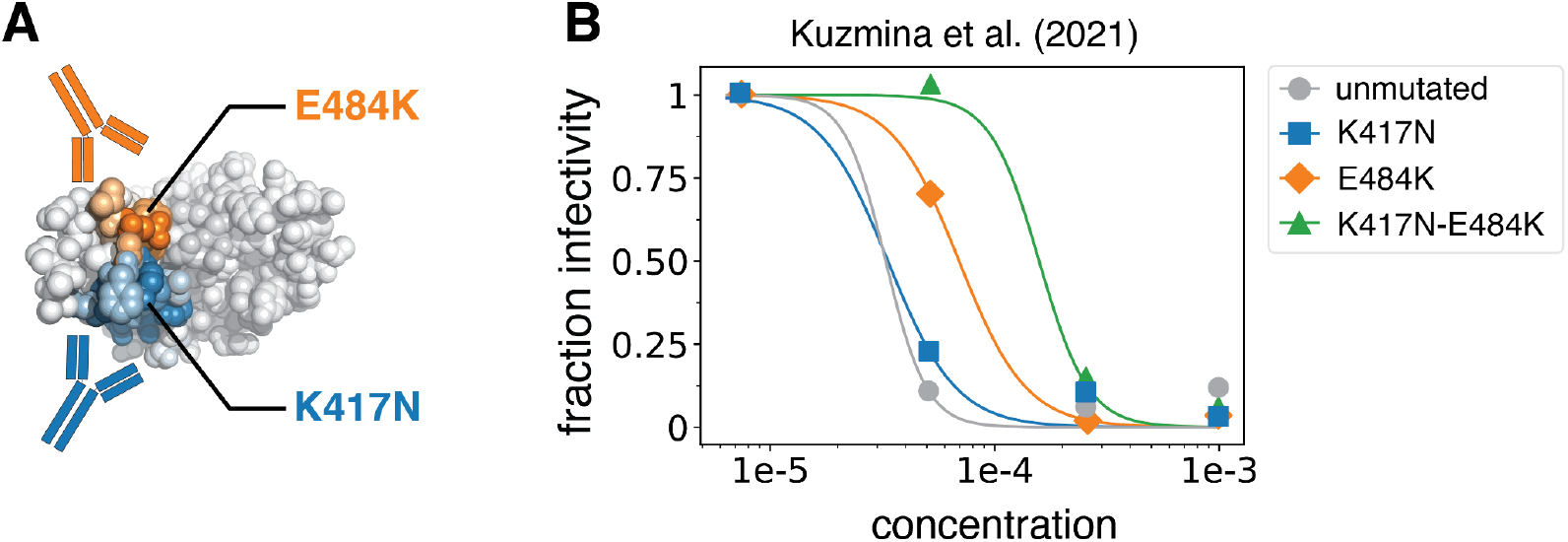
Mutations at multiple epitopes have a synergistic effect on viral escape in real experimental data. **A)** Structure of the SARS-CoV-2 RBD highlighting locations of the K417N and E484K mutations in the class 1 and 2 epitopes, respectively. **B)** Neutralization of unmutated SARS-CoV-2 spike, the K417N and E484K single mutants, and the double mutant against pooled serum from vaccinated individuals. Note how K417N alone has little effect on neutralization, but does cause substantial additional escape in the background of E484K. Experimental data was taken from Kuzmina et al. (Kuzmina *et al*. 2021) and replotted here.

### Biophysical modeling of polyclonal antibody escape

Here we formalize the epitope-based model of polyclonal antibody escape described above in terms of experimental measurables and relevant biophysical quantities. First, let *p*(*v, c*) be the fraction of viral variant *v* that escapes a mixture of polyclonal antibodies at concentration *c*. The quantity *p* (*v, c*) is an experimental measurable, for instance from a neutralization assay. Additionally, suppose the antibodies can bind to one of *E* epitopes on the viral antigen. Let *U_e_* (*v, c*) be the fraction of variant *v* that have an unbound epitope *e* when the antibody mixture is at concentration *c*. Assuming antibodies bind independently without competition, we can express *p*(*v, c*) as:

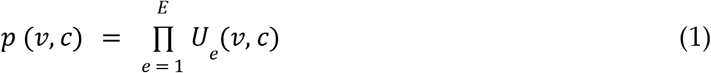

Note that relaxing the assumption that antibodies bind independently without competition to each epitope has little effect on *p*(*v, c*) (see **Appendix**).

Next, we can define *U_e_* (*v, c*) in terms of underlying biophysical properties. Again assuming that there is no competition among antibodies binding to different epitopes, and assuming that the antibody binding to a given epitope *e* can be described by a Hill curve with coefficient of one, then *U_e_* (*v, c*) is given by a Hill equation (Einav and Bloom 2020):

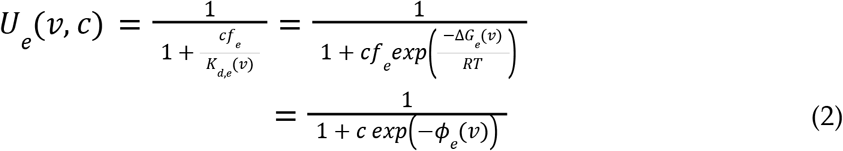

where –*ϕ*_*e*_ (*v*) represents the functional activity of antibodies to epitope *e* against variant *v*. Note that *ϕ*_*e*_ (*v*) depends on both antibody-intrinsic factors (i.e., affinity as quantified by the free energy of binding Δ*G*_*e*_ (*v*)) and extrinsic factors (i.e., relative fraction *f*_*e*_ of antibodies in the mix that bind epitope *e*). The overall functional activity directed to an epitope is a combination of these intrinsic and extrinsic factors, and can be expressed as 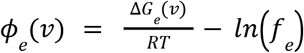. Note that *RT* is the product of the molar gas constant and the temperature, and 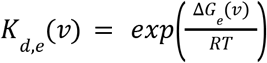 is the dissociation constant. Larger values of –*ϕ*_*e*_ (*v*) indicate that antibodies that bind epitope *e* have a higher functional activity against variant *v*.

Lastly, we can write *ϕ*_*e*_ (*v*) in terms of the contributions of specific mutations. To do this, assume that mutations have additive effects on the free energy of binding for antibodies targeting any given epitope *e* (Otwinowski 2018; Otwinowski, McCandlish and Plotkin 2018). Specifically, let *a*_*wt,e*_ be the functional activity of antibodies that bind epitope *e* on the unmutated viral antigen, with larger values of *a*_*wt,e*_ indicating stronger antibody activity targeting this epitope. Let *β*_*m,e*_ be the extent to which mutation *m* reduces antibody binding or neutralization at epitope *e*, with larger values of *β*_*m,e*_ corresponding to a larger contribution to antibody escape. So, *ϕ_e_* (*v*) can also be written as:

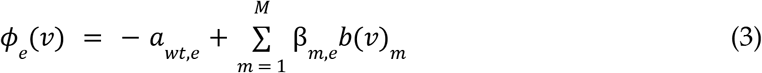

where *b*(*v*)_*m*_ is one if variant *v* has mutation *m* and 0 otherwise, and *m* ranges over all *M* possible mutations. Taken together, the above equations relate the pre-mutation functional activity of antibodies at each epitope (*a*_*wt,e*_) and the antibody escape effects of individual mutations at each epitope (*β*_*m,e*_) to the experimentally measured fraction *p*(*v, c*) of variants *v* that escape an antibody mixture at concentration *c*.

### Fitting biophysical models to deep mutational scanning data

We reasoned that the parameters of our biophysical model could be learned from experiments that measure *p*(*v, c*) for a large number of variants, containing most possible single mutants and a sizable number of multiple mutants (Fowler *et al*. 2010; Kinney *et al*. 2010). These rich genotype-phenotype measurements can be obtained by viral deep mutational scanning. Briefly, viral deep mutational scanning involves generating a large library of viral or viral protein variants that contain single or multiple amino acid mutations, incubating this library with antibodies or sera, and using deep sequencing to identify which variants successfully escape binding or neutralization (Dingens *et al*. 2017; Lee *et al*. 2019; Wu *et al*. 2020; Greaney *et al*. 2021a, 2021c). Although this approach can only measure an infinitesimal fraction of all combinations of mutations, it does measure *p* (*v, c*) for each mutation in many different backgrounds, which should be sufficient for revealing the epitopes targeted by polyclonal antibody mixtures and their associated escape mutations.

To fit our biophysical model using deep mutational scanning data, we created a Python software package named *polyclonal* that uses gradient-based optimization to fit the model to a large set of viral variants *v* and their corresponding experimentally measured escape values, *p*(*v, c*). This software package estimates the *a*_*wt,e*_ and *β*_*m,e*_ parameters that best predict the measured *p*(*v, c*) under tunable and biologically motivated constraints. We typically enforce a sparsity constraint that encourages most *β*_*m,e*_ values to be close to 0, as most mutations to a viral antigen should not mediate antibody escape. We also usually enforce an evenness constraint, as most mutations to a site that mediates antibody escape at a specific epitope tend to have similar effects (or *β*_*m,e*_’s). The software itself is available at https://github.com/jbloomlab/polyclonal and detailed documentation is at https://jbloomlab.github.io/polyclonal. The software also provides methods to visualize the resulting mutation-level escape values in interactive plots as described in the documentation.

### Validation with computationally simulated deep mutational scanning data

To validate the performance of our software, we simulated a hypothetical polyclonal antibody mixture that contains antibodies targeting three major neutralizing epitopes (class 1, 2, and 3) on the SARS-CoV-2 RBD (**Fig. 4A**) (Barnes *et al*. 2020; Greaney *et al*. 2021b). We assigned each epitope a different pre-mutation functional activity (*a*_*wt,e*_) based on the order of typical neutralizing activities against these epitopes in actual human polyclonal serum (class 2 > class 3 > class 1) (Greaney *et al*. 2021b) (**Fig. 4B**). Next, we assigned *β*_*m,e*_ values using prior deep mutational scanning studies (Starr *et al*. 2021b, 2021c) that measured the effects of all single RBD mutations on escape from prototypical monoclonal antibodies targeting each epitope (**Fig. 4C**). Specifically, we used LY-CoV016 (etesevimab) for the prototype class 1 antibody, LY-CoV555 (bamlanivimab) for class 2, and REGN10987 (imdevimab) for class 3 (Starr *et al*. 2021b, 2021c). Note the site of greatest total escape (the sum of positive *β*_*m,e*_ values at a site) is 417 for class 1, 484 for class 2, and 444 for class 3.

**Figure 4.**
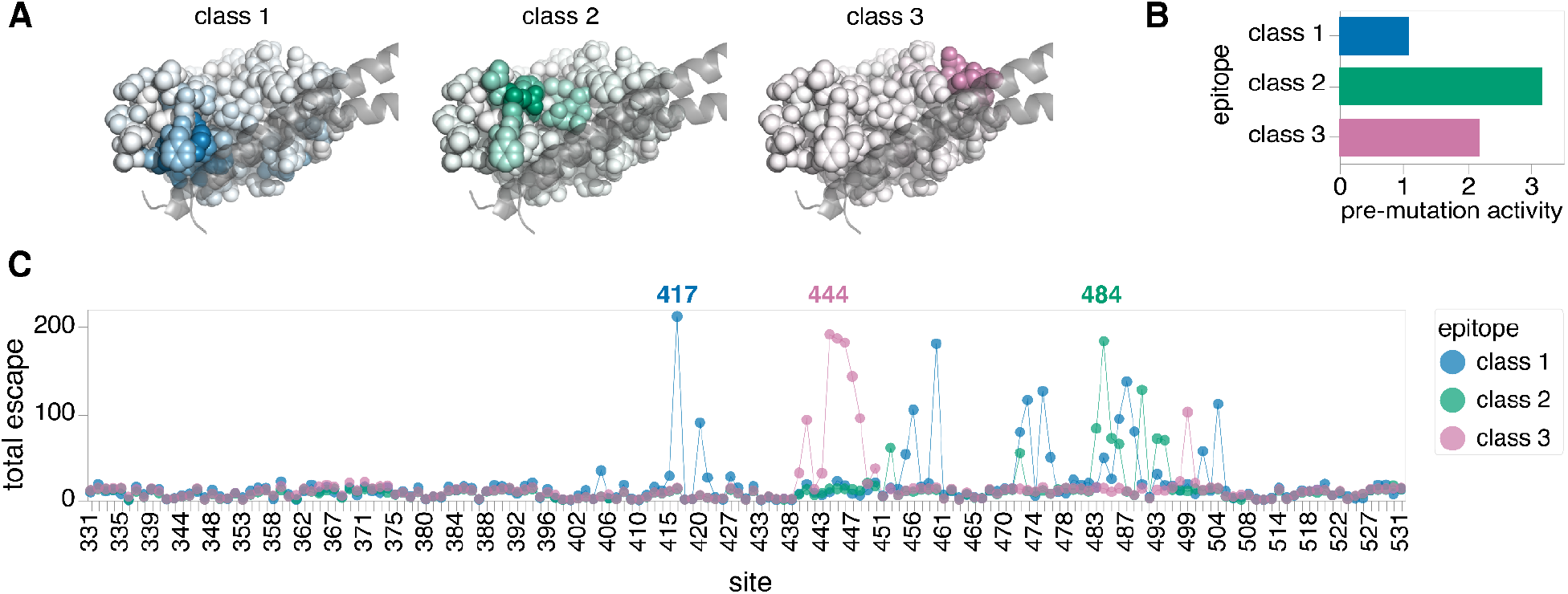
A hypothetical polyclonal antibody mixture targeting the SARS-CoV-2 RBD. **A)** Structures of the SARS-CoV-2 RBD in complex with key helices of ACE2. RBD sites are colored to indicate the extent to which they mediate escape from antibodies targeting each epitope in the hypothetical polyclonal mixture. **B)** Pre-mutation functional activities of antibodies (a_wt,e_) for each epitope in the mixture. Note the class 2 epitope is immunodominant, whereas the class 1 epitope is subdominant. **C)** Sum of positive mutation escape effects (β_m,e_) at each RBD site for each epitope. An interactive version of a heatmap showing all β_m,e_ values is available at https://jbloomlab.github.io/polyclonal/visualize_RBD.html.

Based on this hypothetical polyclonal antibody mixture, we computationally simulated a realistic deep mutational scanning dataset containing 30,000 RBD variants. These variants contained an average of two amino acid mutations, with the number of mutations per variant following a Poisson distribution (**Supp. Fig. 1A**). These variants also only contained mutations at sites found to be functionally tolerated in prior deep mutational scanning, and these sites were well-represented in the simulated dataset (**Supp. Fig. 1B**). We then calculated the true *p*(*v, c*) for each variant using our biophysical model with the *a*_*wt,e*_ and *β*_*m,e*_ defined above. We did this for three concentrations of the hypothetical serum that represent the IC97.5, IC99.9, and IC99.998 against the unmutated RBD (**Supp. Fig. 1C**). Lastly, we added gaussian noise, *N*(0, 0.05), into the *p*(*v, c*) measurements and then truncated the noisy values to be between 0 and 1 to reflect experimental errors and built in a 1% probability of adding or subtracting a mutation from a variant to portray sequencing errors.

We found that *polyclonal* could successfully infer the underlying properties of our hypothetical polyclonal antibody mixture when fit to the noisy, simulated deep mutational scanning dataset. The predicted *a*_*wt,e*_ values were nearly identical to the true *a*_*wt,e*_ values (**Fig. 5A**), suggesting that we can deconvolve the dominance hierarchy of epitopes targeted by a polyclonal antibody mixture. Furthermore, the predicted *β*_*m,e*_ values strongly correlated with the true *β*_*m,e*_ values (**Fig. 5B**, *R*^*2*^=0.63 for class 1, *R*^*2*^=0.91 for class 2, *R*^*2*^=0.85 for class 3), indicating we can learn the effects of mutations on antibody escape at each epitope. Note the lower correlation for the class 1 epitope can be attributed to its relative subdominance. As shown in **Fig. 2B**, mutations to a subdominant epitope manifest little effect on the functional activity of the overall antibody mixture when the immunodominant epitopes remain unaffected, and so only have measurable effects in the subset of mutants with mutations in more dominant epitopes. Nonetheless, our approach can still learn these subdominant effects reasonably well. Taken together, these results demonstrate that noisy *p*(*v, c*)’s measured by deep mutational scanning can be modeled to reveal fine details of polyclonal antibody mixes: the extent to which antibodies target specific epitopes on a viral antigen and the extent to which mutations escape antibodies to each epitope.

**Figure 5.**
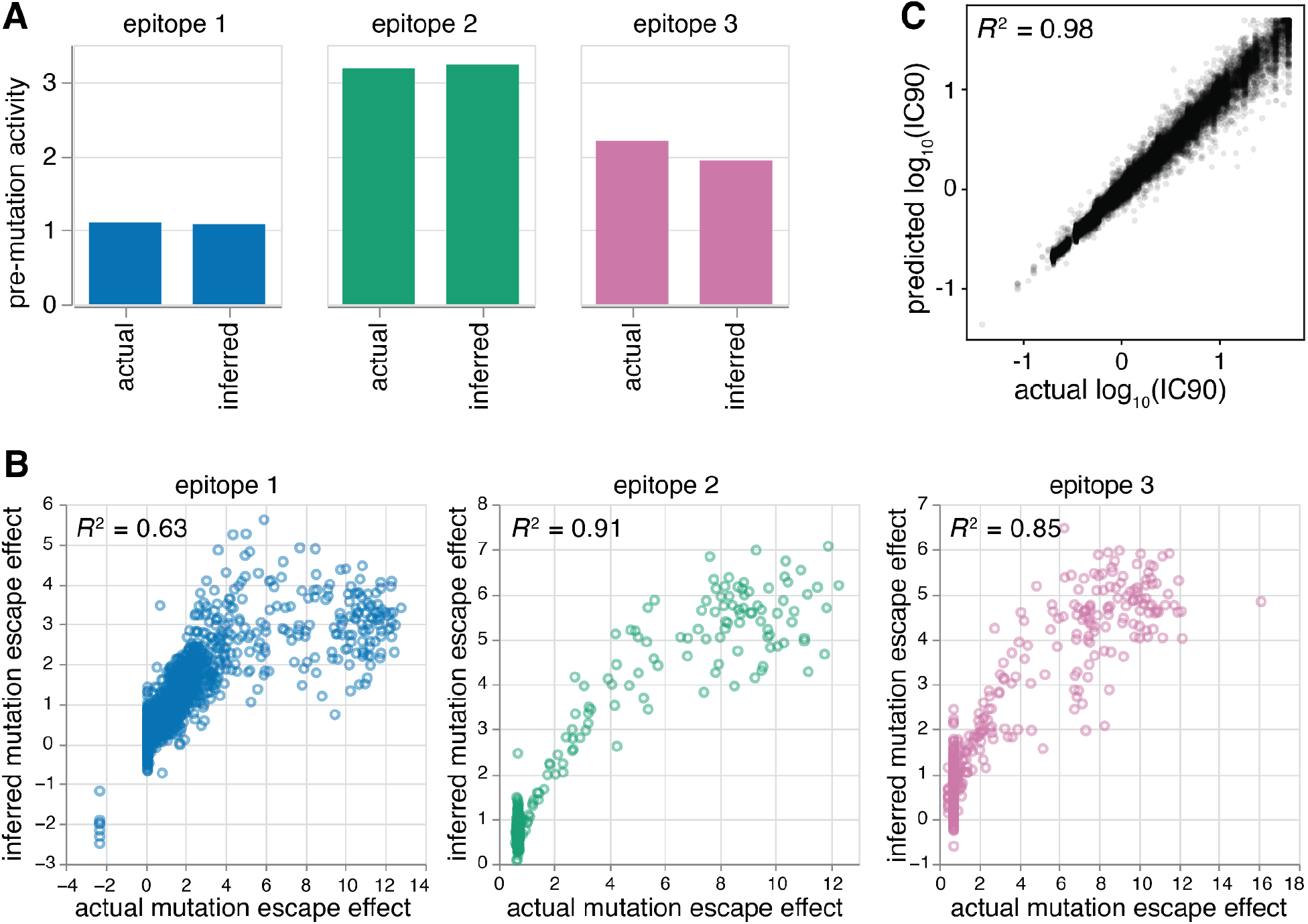
Model validation using a computationally simulated deep mutational scanning dataset. **A)** Pre-mutation functional activities of antibodies (a_wt,e_) inferred by the fit model match the actual a_wt,e_ for each epitope. **B)** Mutation escape effects (β_m,e_) inferred by the fit model strongly correlate with the actual β_m,e_ for each epitope used in the simulation. Mutation escape effects for the immunodominant class 2 epitope are most correlated, while escape effects for the subdominant class 1 epitope are least correlated. **C)** The IC90’s predicted by the fit model strongly correlate with the actual IC90’s of an independently simulated dataset with a higher mutation rate (three mutations on average per variant).

We used the fit biophysical model to predict the extent to which variants that were not seen in our experiments will escape the same polyclonal antibody mixture. To do this, we simulated an independent deep mutational scanning dataset, grounded in the same true *a*_*wt,e*_ and *β*_*m,e*_ values, but with a higher number of mutations (three) on average per variant (**Supp. Fig. 1A**). We found that our fit model could predict the IC90 (the concentration required for *p*(*v, c*) = 0.1) of each variant in the independent dataset with high accuracy (**Fig. 5C**, *R*^*2*^=0.98).

Ultimately, *polyclonal* can be applied to make predictions over all variants with mutations that were observed by deep mutational scanning.

### Model fitting is dependent on experimental design

We next sought to clarify the experimental conditions for deep mutational scanning that lead to accurate model fitting with *polyclonal*. To that end, we systematically explored the impact of three important experimental conditions on model fitting through simulation (see **Methods**).

We first examined the effect of library mutation rate, an important consideration for library design because variants containing multiple mutations are key to revealing epitopes. If a library only contained variants with single mutations, it should not be possible to detect the presence of subdominant epitopes, due to the phenomenon in **Fig. 2B**. Indeed, we found that a library containing 30,000 variants with an average of one mutation could only infer the *β*_*m,e*_ values for immunodominant class 2 epitope (**Fig. 6A**), highlighting the need for including multiply mutated variants. On the other hand, a library containing 30,000 variants with two mutations on average was sufficient for inferring the *β*_*m,e*_ values at all three epitopes, but the accuracy improved as the mutation rate increased (**Fig. 6A**), consistent with our expectation that greater coverage of variants with multiple epitope mutations is helpful. Note that these simulations do not capture the real-world fact that variants will be increasingly less likely to be functional as they accumulate more mutations.

**Figure 6.**
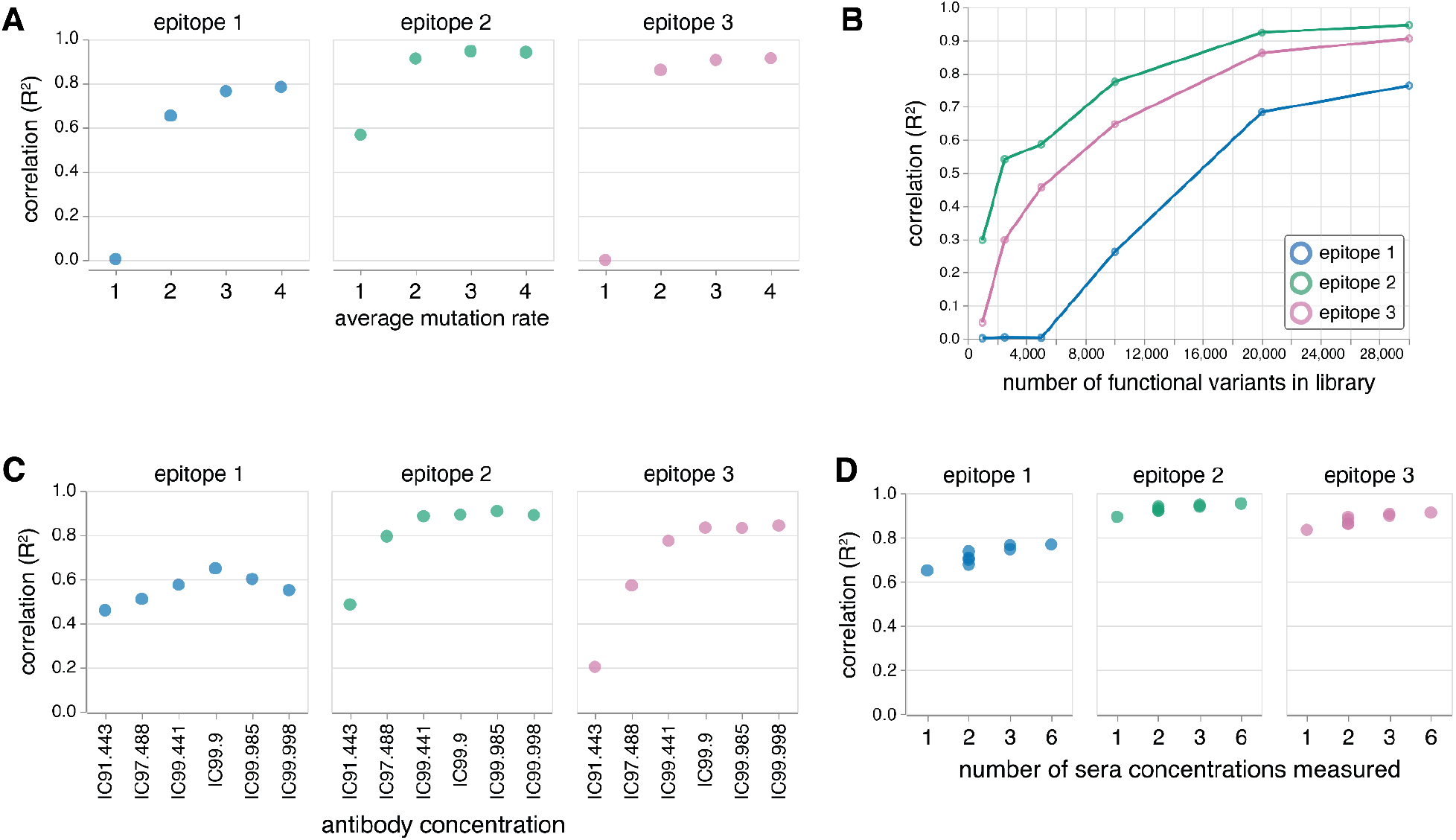
Model fitting on simulated dataset depends on experimental design. **A)** Correlation between inferred and actual mutation escape effects (β_m,e_) improves as the average number of mutations per variant (assuming a Poisson distribution) in the library increases. In particular, the subdominant epitopes (epitopes 1 and 3) can only be accurately fit in libraries with mutation rates higher than one. **B)** Correlation between inferred and actual mutation escape effects (β_m,e_) improves as the number of functional variants in the library increases. **C)** Correlation between inferred and actual mutation escape effects (β_m,e_) depends on the concentration of the antibody mixture. **D)** Correlation between inferred and actual mutation escape effects (β_m,e_) improves as the number of antibody concentrations measured and used to fit the model increases. For details on concentrations used, see: https://jbloomlab.github.io/polyclonal/concentration_set.html.

Next, we investigated the effect of library size, another important library design consideration. If there are too few variants in the library, mutations may not be observed in enough backgrounds to be able to deconvolve their effects on different epitopes. Above we showed that a simulated library with 30,000 variants with three mutations on average could accurately infer the *β*_*m,e*_ values at all three epitopes. Therefore, we set out to determine the minimum number of variants, with three mutations on average, required to accurately infer the *β*_*m,e*_ values at all three epitopes. We found that at least 20,000 functional variants were required to accurately infer the *β*_*m,e*_ values at each epitope (**Fig. 6B**). We also noticed the *β*_*m,e*_ values of immunodominant epitopes can be inferred with fewer variants (**Fig. 6B**), consistent with the idea that effects of mutations at subdominant epitopes can only be observed in the backgrounds where the immunodominant epitope is already mutated.

Lastly, we explored the effect of antibody concentration, an important consideration for deep mutational scanning selections. If the concentration is too high, all variants will be neutralized. If the concentration is too low, there will be little measurable effect from the mutations. To test the effect of different concentrations, we again used the simulated library containing 30,000 variants with three mutations on average. First, we tested if data collected at a single concentration was sufficient. We found that a single concentration of IC99.9 against the unmutated RBD was most effective at inferring the *β*_*m,e*_ values for each epitope, but there does indeed exist a fine balance (**Fig. 6C**). Particularly for the subdominant class 1 epitope, the accuracy is lower when the concentration is too low or too high. We then tested if model fitting could be improved by including additional concentrations flanking the IC99.9, spanning the IC91.443 to IC99.998. Indeed, we found that the accuracy increases as the number of concentrations measured increases (**Fig. 6D**), consistent with our expectation that obtaining more data points along the binding curve is helpful for quantifying the effects of mutations.

## Discussion

We have described a biophysical model of viral escape from polyclonal antibodies. Notably, this model is composed of easily interpretable parameters that capture the interactions between antibodies and the viral epitopes they bind. In addition, we developed a software package that can infer these parameters using deep mutational scanning data. Using a simulated deep mutational scanning dataset, we demonstrated that this approach can infer true parameters from noisy experiments if the deep mutational scanning experiment is appropriately designed.

There are several limitations to our approach. Grouping antibodies by discrete epitopes is an approximation because each antibody is unique, however, decades of experimental work has shown that an epitope-based representation offers a useful way to interpret viral escape. Our model also does not explicitly consider the fact that there are multiple antigens per virion, and instead models escape as simply being the state where no antibodies bind a single idealized antigen. Additionally, our model assumes a specific shape of antigenic epistasis: mutations have additive effects on binding affinity at each epitope, but can manifest non-linear effects on the overall measured phenotype (e.g., neutralization) due to both the non-linear Hill curves that relate binding affinity to total fraction bound at an epitope and the product of these fractions across epitopes. Note that the single-epitope version of our model is similar to the global epistasis models that have proven so useful for interpreting some other types of deep mutational scanning data (Otwinowski, McCandlish and Plotkin 2018; Tareen *et al*. 2022). Our model also assumes that antibody binding to one epitope does not influence the affinity of antibodies to other epitopes, and that the Hill curves have a coefficient of one. It is possible that these assumptions could be relaxed in elaborated versions of our model. We expect that these assumptions will hold reasonably well for viral variants with modest numbers of mutations, but may not extrapolate accurately to variants with dozens of mutations relative to the parental strain used for the deep mutational scanning experiment.

Despite these limitations, our model not only predicts the escape potential of viral variants with arbitrary combinations of mutations that are observed in a simulated deep mutational scanning library—it also clarifies how escape mutations combine to determine the magnitude of viral escape. While this paper only considers simulated data, we envision that our model can be applied to appropriately designed deep mutational scanning experiments to address two main questions: (1) delineating the epitopes targeted by polyclonal serum, and (2) predicting the antigenic properties of new variants with arbitrary combinations of mutations.

## Methods

### Model fitting

Our goal is to estimate the biophysical model parameters that best predict the deep mutational scanning escape measurements under biologically motivated constraints. We defined a loss function that is robust to outliers:

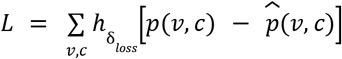

where *p*(*v, c*) is the actual antibody escape fraction for variant *v* at concentration *c*, 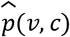 is the predicted antibody escape fraction for variant *v* at concentration *c*, and 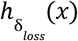 is a scaled Pseudo-Huber function, defined as:

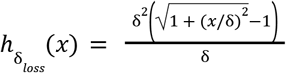

where *δ* is a parameter that indicates when the loss transitions from being quadratic (L2-like) to linear (L1-like). Note that the loss is mostly L2-like when *x*, the residual, is small. However, the loss transitions to become more L1-like when *x* is large, making it more robust to outliers. To minimize this loss function, we use the gradient-based L-BFGS-B method implemented in `scipy.optimize.minimize`.

Additionally, we implemented penalty terms to regularize the parameters to behave under biologically motivated constraints. For detailed information on these penalties and model fitting, see: https://jbloomlab.github.io/polyclonal/optimization.html.

Lastly, fitting the model requires specifying the number of epitopes *a priori*. However, it is not possible to know the number of epitopes that are targeted by antibodies in polyclonal serum in practice. Our approach to resolving this is similar to the “elbow method” commonly used to determine the optimal number of clusters in k-means clustering. We start by fitting a model with one epitope and iteratively fit models with an increasing number of epitopes. At some point, the *N*-th epitope becomes redundant. This is evidenced by a highly negative *a_wt,e_* value (i.e., if antibodies existed against this epitope, they are never bound) and all near-zero *β*_*m,e*_ values, indicating that the previous fit model, containing *N*-1 epitopes, is the one that best describes the polyclonal mixture. To view how this approach was applied to the simulated RBD example, see: https://jbloomlab.github.io/polyclonal/specify_epitopes.html.

### Experimental design simulation

We computationally simulated deep mutational scanning libraries based on the hypothetical polyclonal antibody mixture shown in **Fig. 4**. All libraries contained 30,000 variants, but differed by their mutation rate with variants containing an average of one, two, three, or four mutations. Furthermore, the number of mutations per variant in each library followed a Poisson distribution. Variant escape was also simulated under six concentrations in each library. These concentrations represented the IC91.443, IC97.488, IC99.441, IC99.9, IC99.985, and IC99.998 against the unmutated RBD antigen. For more details on how these experiments were simulated, see: https://jbloomlab.github.io/polyclonal/simulate_RBD.html.

To make comparisons about model fitting, models were fit with identical parameters except for the experimental variable of interest: library mutation rate, library size, and antibody concentration.

To determine the impact of library mutation rate, models were fit to simulated datasets for libraries containing one, two, three, or four mutations on average. All datasets contained 30,000 variants and were measured at three different concentrations.

To determine the impact of library size, models were fit to simulated datasets containing different-sized subsets of variants that were randomly sampled from a library containing an average of three mutations per variant and measured at three different concentrations.

To determine the impact of antibody concentration, models were fit to simulated datasets to a library measured at one or multiple antibody concentrations. This library contained 30,000 variants and three mutations on average per variant.

For more information, see: https://jbloomlab.github.io/polyclonal/expt_design.html.

## Code and Data Availability

Software source code, along with code and data that reproduce the figures (except **Fig. 3**), are available at: https://github.com/jbloomlab/polyclonal

Code and data for reproducing the neutralization curves in **Fig. 3** are available at: https://github.com/jbloomlab/polyclonal-paper

Software documentation is available at: https://jbloomlab.github.io/polyclonal

## Acknowledgements

This work was supported in part by the NIH/NIAID under contract 75N93021C00015 (to JDB), grants R01AI141707 and R01AI165821 (to JDB), and grant F31AI150163 (to WSD). The work was also supported by the NIH/NIGMS CMB Training Grant (T32 GM007270) to TCY and the NSF Graduate Research Fellowship (DGE-2140004) to TCY. JDB and FAM are Investigators of the Howard Hughes Medical Institute.

---

**Supplementary Figure 1.**
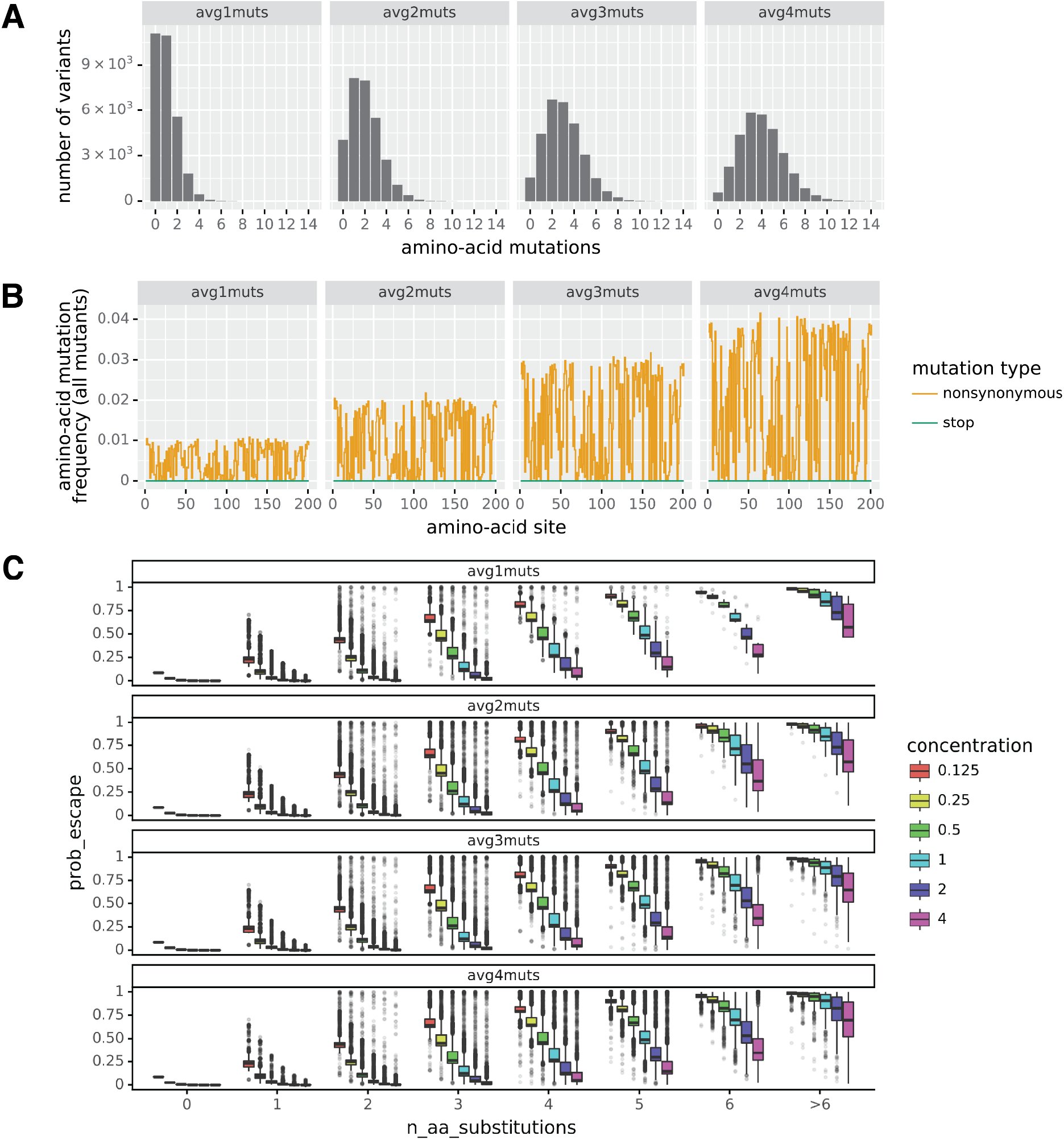
RBD dataset simulation details. **A)** Distribution of the number of amino-acid mutations per variant in each simulated library. **B)** Frequency of mutations at each site on the RBD in each simulated library. **C)** Distribution of antibody escape fractions, *p*(*v, c*), in each simulated library across six concentrations.

## Appendix

We have described a biophysical model that assumes a polyclonal antibody mixture can be divided into independent groups of antibodies that bind to distinct epitopes without competition. Here we interrogate the validity of this assumption, being cognizant of the observation that realistic viral epitopes are often overlapping and therefore not distinct. To do this, we draw from statistical mechanics principles to compare the antibody escape fractions predicted by our independent epitope model and an identically formulated model that instead assumes all epitopes are overlapping.

### 1. Monoclonal antibody case

Before considering the polyclonal antibody case, we first consider the case of a monoclonal antibody that binds a viral antigen. Here, the viral protein can exist in two microstates: bound or unbound by the antibody. The Boltzmann weight is 1 for the unbound state and 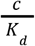 *fo*r the bound state, where *c* is the antibody concentration and *K*_*d*_ is the dissociation constant of antibody-antigen binding. These weights can be derived using the steady-state approximation (Einav and Bloom 2020). We can then define the partition function Ξ as:

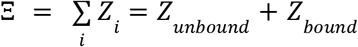

where *Z*_*i*_ represents the Boltzmann weights of the *i m*icrostates. Given this, the probability of a viral antigen being unbound is:

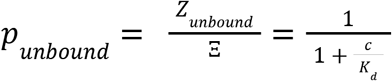

### 2. Polyclonal antibody case

For a polyclonal antibody mixture, we modify *c* to represent the concentration of the polyclonal antibody mixture. We assume that the polyclonal antibody mixture contains antibodies that bind one of *E* epitopes. As follows, the Boltzmann weight of the state where epitope *e* is bound is modified to 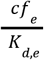,where *f*_*e*_ represents the fraction of antibodies in the mixture that target epitope *e*, and *K*_*d,e*_ is the dissociation constant of antibodies binding to epitope *e*.

#### 2.1 Two distinct epitopes

In a polyclonal antibody mixture, new microstates exist where multiple epitopes are bound by antibodies. For example, we can consider a viral antigen that contains two distinct epitopes (1 and 2) that are targeted by polyclonal antibodies. In addition to the microstates where a single epitope is bound, we now require an additional microstate where both epitopes are bound. The Boltzmann weight for this new microstate is:

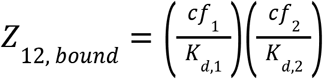

We can then rewrite the partition function Ξ as:

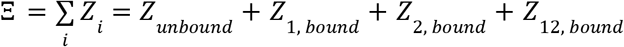

*an*d the probability of a viral antigen being unbound is:

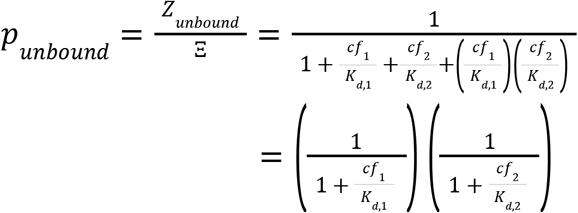

Note that this is the biophysical model that is described in the main text.

#### 2.2 Two overlapping epitopes

In 2.1, the two epitopes were distinct and there was no competition amongst antibodies. However, if the epitopes are overlapping and there is competition, then the microstate where both epitopes are bound (*Z*_12,*bound*_ ) can no longer exist. In this case, the probability of a viral *an*tigen being unbound is:

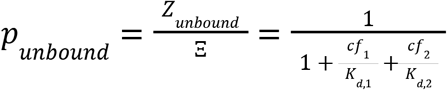

#### 2.3 Extending beyond two epitopes

The same logic applies to viral antigens with more than two epitopes targeted by antibodies. For example, we can write *p*_*unbound*_ for the case of three distinct epitopes as:

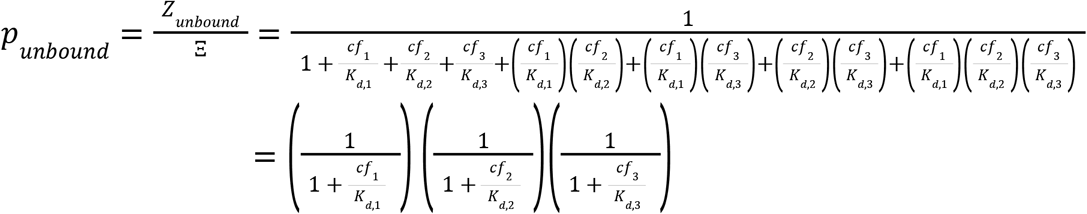

and the case of three overlapping epitopes as:

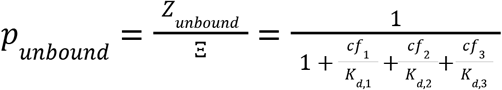

Overall, we found that the predicted *p*_*unbound*_ does not differ by much between the independent and overlapping epitope models under realistic *c*’s, *f*’s, and *K*_*d*_’s. However, the predicted *p*_*unbound*_ becomes more discordant between the two models as the number of epitopes increases, given the number of microstates with multiply bound epitopes increases. In practice, we only fit models to a handful of epitopes and not all of them will overlap. As such, we maintain that the independent epitope assumption is appropriate. To interactively explore how *p*_*unbound*_ differs between the two models, see https://jbloomlab.github.io/polyclonal/partition_function.html.

